# Feral and managed honey bees, *Apis mellifera* (Hymenoptera: Apidae), in southern California have similar levels of viral pathogens

**DOI:** 10.1101/2021.05.17.444546

**Authors:** Amy Geffre, Dillon Travis, Joshua Kohn, James Nieh

**Affiliations:** University of California San Diego, Division of Biological Sciences, Section of Ecology, Behavior, and Evolution

**Author notes:** Corresponding author University of California San Diego, 9500 Gilman Dr., MC0116, La Jolla, CA, 92093-0116 USA, 1-858 822 5011.

**Keywords:** pollinator community, feral honey bees, deformed wing virus, acute bee paralysis virus, black queen cell virus

## Abstract

Bees provide critical pollination services but are threatened by multiple stressors, including viral pathogens. Most studies of pollinator health focus on managed honey bees (*Apis mellifera Linnaeus*) (MHB) or native bee species, but a third player, the feral honey bee (FHB), requires further study. Spillover and spillback of viral pathogens between these managed, feral, and native bees is generating increasing interest. In this case study, we provide evidence suggesting that FHB colonies play an important role in viral pathogen dynamics of southern California pollinator communities because they act as reservoirs, of viral pathogens such as acute bee paralysis virus (ABPV), black queen cell virus (BQCV), and deformed wing virus (DWV). Surprisingly, even though FHB are not treated for diseases or parasites, they harbor similar pathogen loads to MHB, which are usually highly treated, suggesting the need for future studies to determine if FHB resist or are more resilient to viruses.

## Introduction

Multiple factors, including climate change, poor nutrition, pesticides, parasites, and diseases, affect the health of pollinator communities, including bees (Potts et al. 2010, González-Váro et al. 2013, Vanbergen and The Pollinator Initiative 2013, Manley et al. 2019, Hung et al. 2021). In particular, there is growing evidence that many pathogens can be vectored within and between bee species (Genersch et al. 2006, Fürst et al. 2014, Graystock et al. 2015, Alger et al. 2019a and b). Southern California presents a good system to study this phenomenon because it has a large number of managed honey bee colonies (*Apis mellifera*, MHB), an abundant, widespread, and genetically diverse feral honey bee (FHB) population (Schiff et al. 1994, Kono and Kohn 2015), and one of the most diverse native bee communities globally (Moldenke and Neff 1974, Michener 1979, Frankie et al. 2009).

Because of multiple factors, including densely packed colonies (Lindström et al. 2008, Bordier et al. 2017, Nolan and Deplane 2017), pesticide exposure (Rutter et al. 2019, Harwood and Dolezal 2020), and diminished nutrition (Alaux et al. 2010), MHB face mounting pathogen stress (Mexeiner and Le Conte 2016, Dolezal and Toth 2018, Bartlett et al., 2019 and reviewed in Brosi et al., 2017) that may also be problematic for the larger pollinator community. Honey bee-associated viruses (HBAV), including deformed wing virus (DWV) and black queen cell virus (BQCV) (Furst et al. 2014, Graystock et al. 2015, McMahon et al. 2015, Dolezal et al. 2015), can spill into native bee populations, likely via shared floral resources (Genersch et al. 2006, Levitt et al. 2013, Graystock et al. 2015 and 2016) and that MHB colonies may be important pathogen sinks (Alger et al. 2019a and b). In addition to MHB, southern California supports an abundant population of FHB (Kono and Kohn 1995), making honey bees by far the most common floral visitors in for many abundant plant taxa (Hung et al. 2019). These FHB may also play a role in HBAV transmission and prevalence, and as such, the potential capacity for FHB to act as pathogen reservoirs is a concern for both native bees and MHB given the highly interconnected pollinator community in southern California.

While MHB colonies often require intensive, repeated treatments to mitigate multiple viral pathogens, co-occurring FHB colonies are highly successful throughout southern California without human intervention. The reasons for high FHB abundance are unclear, but may be related to natural selection, which FHB experience in ways that MHB do not. Genetic diversity may play a role because FHB in southern California are admixtures of several honey bee lineages and harbor increased levels of genetic variation relative to MHB (Schneider et al. 2004, Kono and Kohn 2015, Cridland et al. 2018, Lin et al. 2018, Calfee et al. 2020). Other studies suggest that distinctive behavioral (Schneider et al. 2004, Loftus et al. 2016, Herb et al. 2018) and physiological traits (Thaduri et al., 2019; Hinshaw et al. 2021) among FHB may increase their ability to deal with pathogens.

In this case study, we take a key first step to understanding the connections between HBAV in FHB and MHB in southern California by measuring the abundance of important viral pathogens. We assayed MHB and FHB colonies for two common HBAV known to infect other pollinator taxa, deformed wing virus (DWV), and black queen cell virus (BQCV) (Genersch et al. 2006, Greystock et al. 2015, Tehel et al. 2016, Alger et al. 2019a and b), as well as another important apiary virus, acute bee paralysis virus (ABPV), seven times over a year.

## Methods

### Feral and native bee collections

We collected samples from 3 FHB and 3 MHB colonies from San Diego and southern Riverside counties (Figure 1) during September, October, November of 2019, and April, June, July, and August of 2020. We selected MHB colonies that were managed according to California beekeeping standards, with active queens from commercial queen breeders. To ensure that we collected feral bees, rather than recent swarms from managed apiaries, we sampled FHB colonies which had been recorded at that location for at least a year. Colonies were located at least 8 km apart to minimize overlap or drifting of foraging bees between colonies. Our study originally included an additional MHB apiary and 4 additional FHB colonies, but these were lost due to limited access to sites and drop-out of commercial beekeepers during the COVID-19 pandemic and natural attrition.

**Figure 1.**
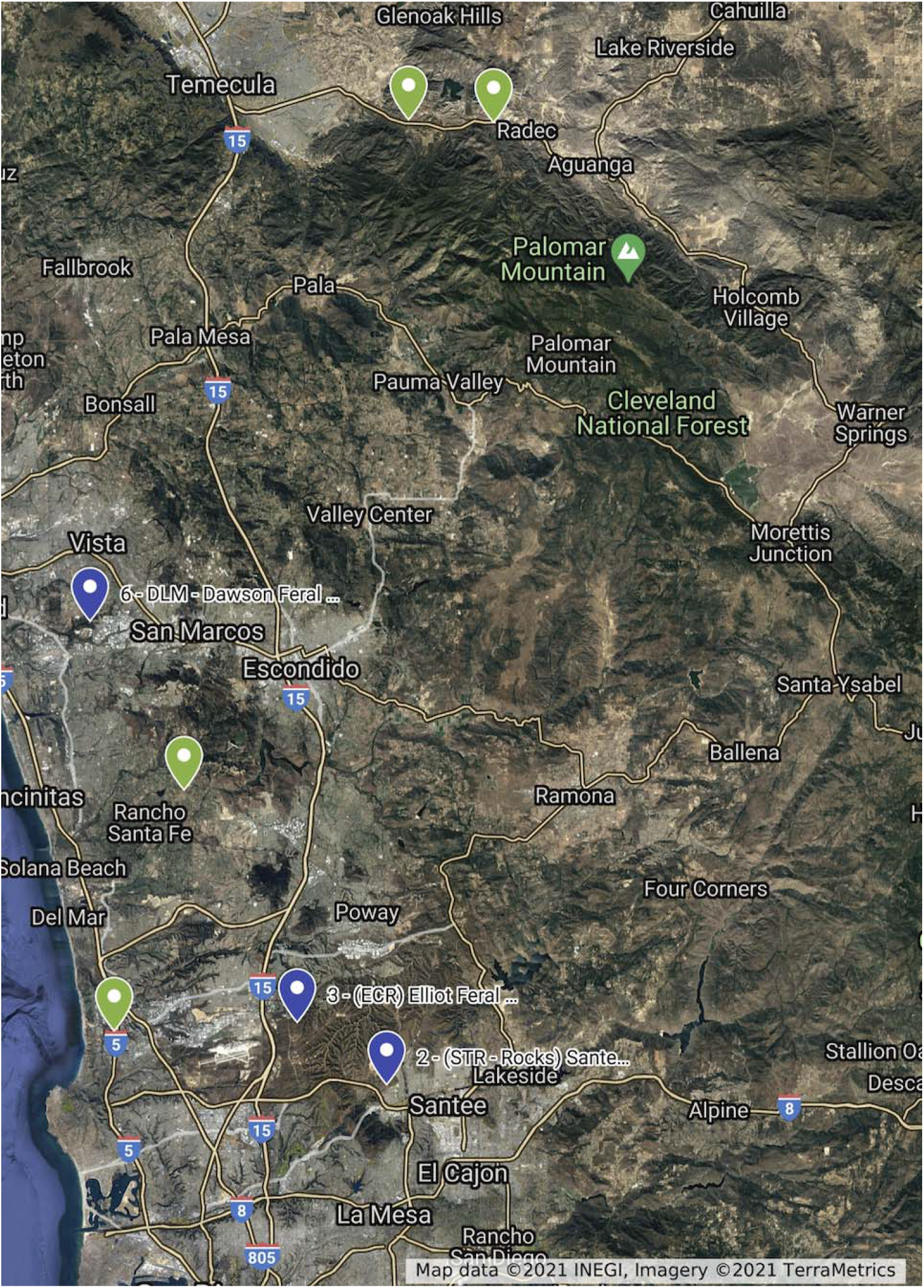
Collection sites (purple markers = feral colonies, green markers = managed colonies). Image from Google Maps.

**Figure 2:**
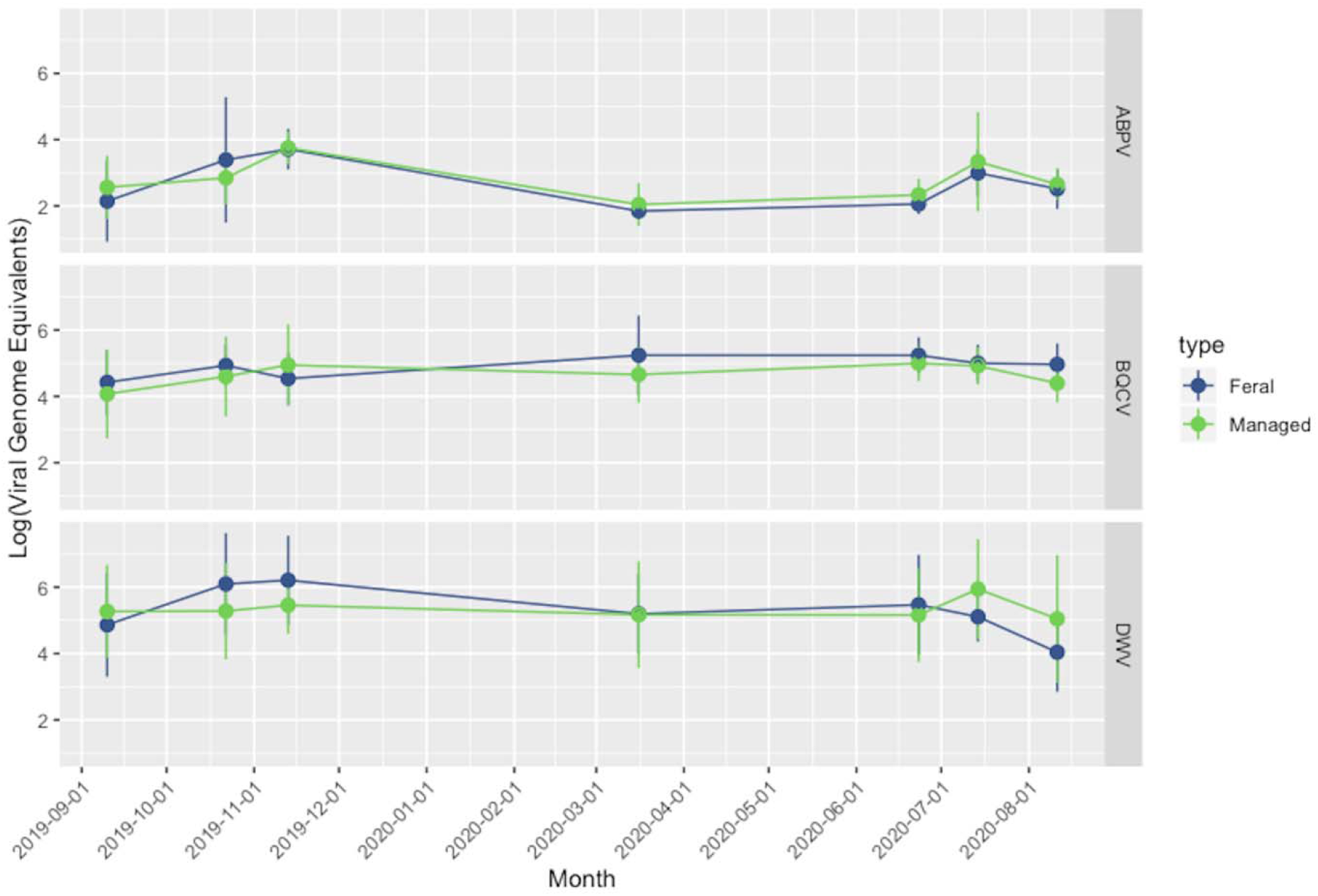
Infection intensity (quantified as the log of copies of viral genome equivalents (VGE) per 100 ng of extracted RNA per honey bee) of feral (purple) and managed (green) honey bees across seven months. Error bars indicate standard deviation. At each month (time point) the sample size is 5 bees per colony (3 FHB and 3 MHB colonies), except for November, when the sample size was 3 bees per colony. Winter months (December through March) were omitted because foragers were difficult to collect due to low temperatures (ABPV = Acute bee paralysis virus; BQCV = Black queen cell virus; DWV = Deformed wing virus). The proposed cut-off for infection viability is Log (Viral Genome Equivalents) ~ 2, based upon Dolezal et al., 2016).

We collected foragers since these could be obtained from entrances of FHB and MHB colonies, without altering the colony, and because foragers can transmit viruses between colonies during robbing or drifting and via shared floral resources. Bees were only collected on sunny or partially sunny days with little wind to ensure natural foraging activity. Bees from each colony on any given day were collected into a single 15 mL centrifuge tube and immediately humanely euthanized on dry ice for later HBAV characterization by qPCR. All samples were stored at −70°F until RNA extraction. To avoid cross-contamination between sites, we used disposable gloves while collecting bees, and disinfected nets with 10% bleach for at least 10 min between sites.

All MHB were sampled with the permission of apiary owners. Locations and identities of participating apiaries were kept confidential. FHB sites were either on public land or University of California Natural Reserve Systems sites. While individual bees were collected, we did not otherwise alter the colonies. No protected or endangered species were involved in this survey.

### RNA Extractions and Quantitative PCR

Total RNA was extracted from whole individual bees using Trizol™ (Life Technologies), according to an established protocol for quantifying HBAV titers (Carillo-Tripp et al. 2016). Afterwards, all samples were resuspended in HyClone molecular-biology-grade water (Cytiva). Sample concentration and quality were assayed with a Nanodrop 2000 spectrophotometer (Thermofisher). Both 260/230 and 260/280 values were required to be 2.00 (+/− 0.20) for qPCR. All sample concentrations were equalized to 100ng/uL (+/− 20ng/uL).

We quantified absolute virus titers in each sample for DWV, BQCV, and ABPV with primers published in Carillo-Tripp et al. (2015). Samples were quantified using one-step qRT-PCR (iTaq™ Universal SYBR^®^ Green One-Step Kit (BioRad)) on the CFX Maestro platform (BioRad) according to manufacturer’s guidelines. Approximately 200 ng (2 uL) of each sample was used per well. All samples were run in duplicate. The absolute number of viral genome equivalents present in each bee was determined by comparing the sample Cq with a serial dilution of a known standard curve derived from HBAV fragments linearized in a plasmid, and constructed at University of Illinois, Urbana-Champaign (Carillo-Tripp et al. 2016).

### Statistical Analyses

Viral titers for sites and bee types (FHB or MHB) were compared using repeated measures general linear models (GLM). These analyses were carried out in RStudio (R Core Team 2019, RStudio Team 2020), using the lme4 (Bates et al. 2015), nlme (Pinhero et al. 2019) and lmerTest (Kuznetzova et al. 2017) packages. In our model, bee type and date of collection were set as primary effects; when site showed a significant effect on virus titer (in GLMs within bee types for each virus), it was included as a random effect. To normalize viral titer data, we log-transformed viral genome equivalents (log(VGE)) to improve model fit. All graphics were produced using ggplot2 (Wickham, 2016). All raw data and analysis code is available via GitHub upon publication.

## Results

Average viral titers in our colonies fluctuated over the collection dates (see Table 1 for complete statistics), as is typical among honey bee colonies (Antúnez et al. 2015), with the exception of ABPV, which was present only in low titers, often below Log (VGE) ~ 2, which is the minimum level that likely represents true infection (Dolezal et al. 2016). However, viral titers of FHB and MHB did not differ significantly during any sampling period, although DWV trended towards a difference between bee types, over time (see Table 1 for complete statistics, and Figure 1).

**Table 1.** Repeated measures regression of viral titers in FHB and MHB over sampling dates. Model: log (*Viral genome equivalents of target virus*) ~ *Bee type* + *Date* + (*Bee type* × *Date*) + (1|*Site*), where *Bee type* and *Date* are fixed effects, and *Site* is a random effect.

| **ABPV** | **Effect Size Estimate ($\beta$)** | **Standard Error** | **DF** | **t-value** | **p-value** |
| --- | --- | --- | --- | --- | --- |
| Intercept | 24.120 | 12.980 | 248.400 | 1.858 | 0.064 |
| *Bee Type* | -20.230 | 18.250 | 248.400 | -1.108 | 0.269 |
| *Date* | -0.001 | 0.001 | 248.400 | -1.654 | 0.099 |
| *Type x Date* | 0.001 | 0.001 | 248.300 | 1.114 | 0.267 |

| **BQCV** | **Effect Size Estimate ($\beta$)** | **Standard Error** | **DF** | **t-value** | **p-value** |
| --- | --- | --- | --- | --- | --- |
| Intercept | -22.840 | 10.010 | 248.500 | -2.281 | 0.023 |
| *Bee Type* | 3.635 | 14.070 | 248.500 | 0.258 | 0.796 |
| *Date* | 0.002 | 0.001 | 248.100 | 2.773 | 0.006 |
| *Type x Date* | 0.000 | 0.001 | 248.100 | -0.281 | 0.779 |

| **DWV** | **Effect Size Estimate ($\beta$)** | **Standard Error** | **DF** | **t-value** | **p-value** |
| --- | --- | --- | --- | --- | --- |
| Intercept | 44.250 | 16.780 | 248.400 | 2.637 | 0.009 |
| *Bee Type* | -44.010 | 23.610 | 248.400 | -1.864 | 0.064 |
| *Date* | -0.002 | 0.001 | 248.400 | -2.326 | 0.021 |
| *Type x Date* | 0.002 | 0.001 | 248.400 | 1.869 | 0.063 |

## Discussion

FHB and MHB had similar levels of HBAV infection in southern California, suggesting that both bee types may transmit viruses to the wider bee community. Recently, Hinshaw et al. (2021) working in Pennsylvania found similar (and at times higher levels of HBAV) in FHB as compared to MHB and other studies have found several viruses present in similar or higher titers in FHB, compared to MHB (in Georgia, Bartlett et al. 2021, and in the UK, Thompson et al., 2014). The fact that FHB across North America appear to be reservoirs of viruses that negatively affect MHB, and perhaps other pollinators, means their role cannot be ignored when considering the dynamics of viral spread in pollinator communities.

FHB apparently thrive despite constant exposure to the same viruses that require treatment in MHB colonies. Natural selection in FHB populations seems to have increased their ability to ward off severely negative consequences of HBAV (Seeley 2007; Thaduri et al., 2019). In southern California, the increased genetic diversity of FHB populations may allow them to better deal with HBAV [as suggested by previous work in other bee subpopulations observing various immune stressors (Tarpy 2003, Meixner et al. 2010, Youngstead et al., 2015; Mikheyev et al. 2015)] and an important HBAV vector (Moretto et al. 1999). However, increased genetic diversity may not be required for adaptation against common diseases. In South Carolina (López-Uribe et al. 2017), FHB have somewhat lower genetic diversity than MHB, though expression levels of immune response genes are positively correlated with the genetic diversity of FHB colonies. In southern California, common FHB behaviors such as increased swarming frequency, smaller colony sizes, and enhanced defensive behavior that may help ward off diseases are often attributed to admixture with the African subspecies, *A. mellifera scutellata* (Schneider et al. 2004, Loftus et al. 2016, Herb et al. 2018, Carr et al. 2020). Still, in other locations without such ancestry, similar behaviors can occur (Seeley et al. 2015 and 2017, Locke 2016, Loftus et al. 2016) suggesting that feralization and natural selection may promote such beneficial behaviors even in the absence of *A. mellifera scutellata* ancestry.

Given the multiple factors currently threatening pollinators, especially bees, (Potts et al. 2010, González-Váro et al. 2013, Vanbergen and The Pollinator Initiative 2013, Manley et al. 2019, Hung et al. 2021) there is growing need to find suitable ways to mitigate pathogen pressure among our important pollinators. To protect our critical managed bees (Mexeiner and Le Conte 2016, Dolezal and Toth 2018, Rutter et al. 2019), and diverse bee communities (Genersch et al. 2006, Fürst et al. 2014, Graystock et al. 2015, Alger et al. 2019a and b), there is strong impetus to find new and sustainable means to decrease honey bee HBAV loads. Current knowledge of FHB suggests there may be untapped resources to mitigate viral pathogen stress. More studies are needed, but our results suggest that the physiological and behavioral mechanisms used by FHB to deal with HBAV should be further explored. We hope our case study will motivate research into the role of FHB in pathogen dynamics of pollinator communities and into whether and how they ward off serious infection despite exposure to HBAV that seriously impact MHB.

## Acknowledgements

We thank community beekeepers in San Diego and Riverside counties for providing managed bees, anonymous reviewers and members of Nieh and Kohn Labs for valuable feedback, Dolezal Lab at UIUC for providing the USR viral standard curve, and the UC Academic Senate, UCSD Jeanne M. Messier Endowment Fund and UCNRS Mildred E. Mathias Graduate Student Research Grant Program for funding.

