## Supplementary material for "Feral and managed honey bees, *Apis mellifera* (Hymenoptera: Apidae), in southern California have similar levels of viral pathogens": Data and R Code: GeffreEtAl-2021_RCode.Rmd

````
---
title: "Feral Bee Pathogen Survey - Monthly Comparisons with Corrected Values"
output:
  html_document:
    toc: true
    toc_float: true
---

```{r setup, include=FALSE}
knitr::opts_chunk$set(echo = TRUE)
```

# Feral Honey Bee Pathogen Survey - Monthly Comparisons

## Introduction 

The prevalence of honey-bee associated viruses is typically only assayed among managed honey bees.  To better understand the dynamics of these viruses in the pollinator community, we suveyed the viral loads of a suite of feral honey bee colonies living intermingled with managed colonies. We hope to use these data to better understand how this group of typically understudied bees contributes to pathogen dynamics in the pollinator communities they live in.

This material is to be associated with the paper "Feral and managed honey bees, Apis mellifera (Hymenoptera: Apidae), in southern California have similar levels of viral pathogens", by Amy Geffre, Dillon Travis, Joshua Kohn and James Nieh (UCSD, Biological Sciences: Ecology, Behavior and Evolution).

## Data Details

### Biological Details
For each sample, background data should include: 

1. Sample ID ("sample")
2. Site
    + Latitude and longitude ("lat" and "long")
    + Acronym ID ("site")
4. Bee type ("type")
    + Feral
    + Managed
5. Month
    + "date" - a date value of sampling month
    + "month2" - a numeric value of sampling month
    + "month3" - a factor variable of sampling month

We will consider bee type and month as fixed effects, as we hypothesize virus titers will vary between bee types, over time.   

We will compare titers (as "Starting Quantity" or SQ) of four notable bee viruses, as calculated by absolute quantification (JCT et al, 2015 - USR method):

* Acute bee paralysis virus (ABPV)
* Black queen cell virus (BQCV)
* Deformed wing virus (DWV)

Each virus will be denoted in the "Target" column, using the acronyms above.

Viruses are defined with the variable "cqmean" (the mean cycle of quantification, between two technical replicates of a sample), "cqstddev" (std. deviation of Cq), "meansq" (mean starting quantity described by absolute quantification), "sq std dev" (standard deviation of the SQ), and "logmeansq"" (the mean SQ logged to improve model fitting).

Please note that, for each month, bees are not sampled on exactly the same date, due to issues of time. However, all bees are sampled within three days of each other for a given month. 

```{r Packages, include=F}
library(ggplot2)
library(ggpubr)
library(dplyr)
library(stats)
library(car)
library(multcomp)
library(MASS)
library(pscl)
library(nlme)
library(lme4)
library(lmerTest)
```

### Analysis Description

I will compare the virus titers over the months by using a repeated measures general linear model. I take the structure for these analysis from [UCLA](https://stats.idre.ucla.edu/r/seminars/repeated-measures-analysis-with-r/) and [M Clark](https://m-clark.github.io/mixed-models-with-R/random_intercepts.html) and [PSU](https://online.stat.psu.edu/stat510/lesson/10/10.1), but modify it for my data.

## Analysis

### Reading in the Data

```{r Data Read-In}
allmonths <- read.csv("GeffreEtAl2021_allmonths.csv")
colnames(allmonths) <- tolower(colnames(allmonths))
allmonths$date <- as.Date(allmonths$date)
allmonths$month2 <- as.numeric(allmonths$month2)
allmonths$month3 <- as.factor(allmonths$month2)
str(allmonths)
allmonths$logmeansq <- log10(allmonths$meansq)
allmonths$logmeansq<-replace(allmonths$logmeansq, allmonths$logmeansq<0,0) 
  # Replace the two -Inf in the results (b/c of 0 values for SQ)
  # This is biologicall functional. 

summary(allmonths)
```

Note here that a few samples (notably several bees from TMM, in fall) have remarkable levels of DWV. In general, the median is normal, though (~5-6 log(VGE)). 

```{r}
all_group_means <- allmonths %>%
  group_by(date, type, target) %>%
  summarise(mean_lms = mean(logmeansq),  # calculates the mean of each group
            sd_lms = sd(logmeansq), # calculates the standard deviation of each group
            n_lms = n(),  # calculates the sample size per group
            SE_lms = sd(logmeansq)/sqrt(n())) # calculates the standard error of each group
```


```{r}
allline <- ggplot(all_group_means, aes(x = date, y = mean_lms, colour = type)) +
  geom_line() +
  facet_grid(rows = vars(target), labeller=labeller(date = allmonths$month)) +
  geom_pointrange(aes
                  (ymin = mean_lms-sd_lms, ymax = mean_lms+sd_lms), 
                  data = all_group_means) +
  geom_point(size = 1) +
  scale_x_date(date_breaks = "1 month") +
  scale_fill_manual(values=c("#39568CFF", "#73D055FF")) +
  scale_color_manual(values=c("#39568CFF", "#73D055FF")) +
  labs(x="Month", y = "Log(Viral Genome Equivalents)") +
  theme(axis.text.x = element_text(angle = 45, hjust = 1), plot.margin = margin(1, 1, 1, 2, "cm"))

allline
```

```{r}
allline2 <- ggplot(all_group_means, aes(x = date, y = mean_lms, colour = type)) +
  geom_line() +
  facet_grid(rows = vars(target), labeller=labeller(date = allmonths$month)) +
  geom_pointrange(aes
                  (ymin = mean_lms-SE_lms, ymax = mean_lms+SE_lms), 
                  data = all_group_means) +
  geom_point(size = 1) +
  scale_x_date(date_breaks = "1 month") +
  scale_fill_manual(values=c("#39568CFF", "#73D055FF")) +
  scale_color_manual(values=c("#39568CFF", "#73D055FF")) +
  labs(x="Month", y = "Log(Viral Genome Equivalents)") +
  theme(axis.text.x = element_text(angle = 45, hjust = 1), plot.margin = margin(1, 1, 1, 2, "cm"))

allline2
```

### Structure Data for Repeated Measures (using Logged Mean(SQ))

```{r Meteing Data into Target Bins ABPV}
# ABPV
abpvall <- allmonths[which(allmonths$target=='ABPV'),]
summary(abpvall)
qqnorm(abpvall$logmeansq)
shapiro.test(abpvall$logmeansq)
```

``` {r BQCV}
# BQCV
bqcvall <- allmonths[which(allmonths$target=='BQCV'),]
summary(bqcvall)
qqnorm(bqcvall$logmeansq)
shapiro.test(bqcvall$logmeansq)
# Normal, according to Shapiro test
```

``` {r DWV}
# DWV
dwvall <- allmonths[which(allmonths$target=='DWV'),]
summary(dwvall)
qqnorm(dwvall$logmeansq)
shapiro.test(dwvall$logmeansq)
```


### Within Beetypes

```{r Meteing Data into Type Bins}
# FHB
feralall <- allmonths[which(allmonths$type=='Feral'),]
summary(feralall)

# MHB
manageall <- allmonths[which(allmonths$type=='Managed'),]
summary(manageall)

```

```{r}
# ABPV
abpvferal <- feralall[which(feralall$target=='ABPV'),]
summary(abpvferal)
qqnorm(abpvferal$logmeansq)
shapiro.test(abpvferal$logmeansq)
```

```{r}
# BQCV
bqcvferal <- feralall[which(feralall$target=='BQCV'),]
summary(bqcvferal)
qqnorm(bqcvferal$logmeansq)
shapiro.test(bqcvferal$logmeansq)
```

```{r}
# DWV
dwvferal <- feralall[which(feralall$target=='DWV'),]
summary(dwvferal)
qqnorm(dwvferal$logmeansq)
shapiro.test(dwvferal$logmeansq)
```

##### LMM for Ferals only
```{r}
feralabpv.glm = glm(logmeansq ~ site*date,
              data=abpvferal)

summary(feralabpv.glm)
# ABPV has a site effect among FHB
```

```{r}
feralbqcv.glm = glm(logmeansq ~ site*date,
              data=bqcvferal)

summary(feralbqcv.glm)
# BQCV does not have a site effect for FHB
```

```{r}
feraldwv.glm = glm(logmeansq ~ site*date,
              data=dwvferal)
summary(feraldwv.glm)
# DWV does not have a site effect for feral bees
```

##### LMM for manages only

```{r}
# ABPV
abpvmanage <- manageall[which(manageall$target=='ABPV'),]
summary(abpvmanage)
qqnorm(abpvmanage$logmeansq)
shapiro.test(abpvmanage$logmeansq)
```

```{r}
# BQCV
bqcvmanage <- manageall[which(manageall$target=='BQCV'),]
summary(bqcvmanage)
qqnorm(bqcvmanage$logmeansq)
shapiro.test(bqcvmanage$logmeansq)
```

```{r}
# DWV
dwvmanage <- manageall[which(manageall$target=='DWV'),]
summary(dwvmanage)
qqnorm(dwvmanage$logmeansq)
shapiro.test(dwvmanage$logmeansq)
```

```{r}
manageabpv.glm = glm(logmeansq ~ site*date,
              data=abpvmanage)

summary(manageabpv.glm)
# Site is important for ABPV titers among MHB
```

```{r}
managebqcv.glm = glm(logmeansq ~ site*date,
              data=bqcvmanage)

summary(managebqcv.glm)
# Site is important among MHB for BQCV
```

```{r}
managedwv.glm = glm(logmeansq ~ site*date, data=dwvmanage)

summary(managedwv.glm)
# No importance of site in DWV titers for MHB
```

### Linear Mixed Effects Model to compare Log(MeanSQ) across year/bee types

To accomodate for skewed data structure, we use a mixed effects model, especially to account for variance introduced by the different sites (though in most cases, we don't see spatial autocorrelation for these viruses). 

Note that lmer produces weird DF: "The parameter estimates calculated by lmer are the maximum likelihood or the REML (residual maximum likelihood) estimates and they are not based on observed and expected mean squares or on error strata.  And that's a good thing because lmer can handle unbalanced designs with multiple nested or fully crossed or partially crossed grouping factors for the random effects." - https://stat.ethz.ch/pipermail/r-help/2006-May/094765.html

#### LMM ABPV
```{r}
abpv.lmer <- lmer(formula = logmeansq ~ type*date + (1|site),
                data = abpvall)
summary(abpv.lmer)
rand(abpv.lmer)
```

```{r}
abpv.lmer1 <- lmer(formula = logmeansq ~ type*month2 + (1|site),
                data = abpvall)
summary(abpv.lmer1)

rand(abpv.lmer1)
```

```{r}
AIC(abpv.lmer, abpv.lmer1)
```


#### LMM BQCV

```{r}
bqcv.lmer <- lmer(formula = logmeansq ~ type*date + (1|site),
                data = bqcvall)
summary(bqcv.lmer)
```

```{r}
bqcv.lmer1 <- lmer(formula = logmeansq ~ type*month2 + (1|site),
                data = bqcvall)
summary(bqcv.lmer1)
```

```{r}
AIC(bqcv.lmer, bqcv.lmer1)
```

```{r}
anova(bqcv.lmer1)
```


#### LMM DWV

```{r}
dwv.lmer <- lmer(formula = logmeansq ~ type*date +  (1|site),
                data = dwvall)
summary(dwv.lmer)
```

```{r}
dwv.lmer1 <- lmer(formula = logmeansq ~ type*month2 +  (1|site),
                data = dwvall)
summary(dwv.lmer1)
```


```{r}
AIC(dwv.lmer, dwv.lmer1)
```
````
